# PRESGENE: A web server for PRediction of ESsential GENE using integrative machine learning strategies

**DOI:** 10.1101/2022.11.25.517801

**Authors:** Sutanu Nandi, Gauri Panditrao, Piyali Ganguli, Ram Rup Sarkar

## Abstract

Study of essential genes in disease-causing organisms has wide application in the prediction of therapeutic targets and exploring different clinical strategies. Predicting gene essentiality for large set of genes in non-model, less explored organisms is challenging. Computational methods that use machine learning (ML)-based strategies are popularly adopted for essential gene prediction as they provide key advantage of considering diverse biological features. Previous works from our group have demonstrated two ML-based pipelines for predicting essential genes with high accuracy that mitigates the problems of sufficient labeled imbalanced dataset and limited labeled datasets of essential genes. Here we present PRESGENE at https://presgene.ncl.res.in, a ML-based web server for prediction of essential genes in unexplored eukaryotic and prokaryotic organisms. Our algorithms mitigate the problems of training dataset imbalance and limited availability of experimentally labeled data for essential genes. PRESGENE with its user-friendly web interface and high accuracy will prove to be a seamless experience for biologists looking for an accurate essential gene prediction server with limited labeled data for novel organisms.

## 1 Introduction

The minimally essential genes in an organism comprise a set of absolutely necessary genes for its survival under any environmental condition(1). The gene essentiality information helps prioritize a set of crucial genes and their functional properties which may serve as important drug targets against various infectious diseases such as Cutaneous and Visceral Leishmaniasis, Tuberculosis, Typhoid, etc. The study of the mammalian essential genes also provides evidence for identifying important therapeutic targets and biomarkers for the treatment of cancer and other diseases. The gene essentiality information of the lesser-studied disease-causing organisms helps to identify and annotate these minimally essential genes that contribute to the understanding of the pathogen biology.

Establishing the essentiality for a large set of genes in non-model, less explored organisms is challenging, as the experimental standardization of protocols for performing genome-wide screens to identify dispensability and sampling for a range of experimental conditions is laborious and time-consuming. Hence, computational techniques based on homology mapping, constraint-based modeling, and machine learning strategies are becoming useful to predict essential genes with high accuracy in a small amount of time(2–4).Machine learning (ML)-based methods offer the key advantage of considering diverse biological features that influence gene essentiality (Table 1).Various data-driven ML-based algorithms have been used for the prediction of essential genes, *e.g*., decision tree(5), random forest(6), logistic regression (5,7), ensemble (5), support vector machine (8–11), probabilistic Bayesian-based methods (5,7,12), K Nearest neighbor (K-NN) and Weighted KNN (WKNN)(13). A major limitation of the existing ML-based methods (Supplementary data, Table S1) is the necessity of large amount of labeled data from experiments and often fails to predict actual essential genes when the labeled data set is imbalanced or insufficient. Moreover, the essential gene prediction web servers built so far are heavily dependent on homology mapping-based strategy alone, while the other biologically relevant features derived from the genome-scale metabolic networks that significantly impact gene essentiality under varied environmental conditions have not been explored sufficiently. Constraint-based modeling strategies, such as Flux Balance Analysis (FBA), employing genome-scale reconstructed metabolic networks, are widely used for predicting essential genes by performing *in-silico* knockout of a gene and estimating its corresponding lethality (14–16). A limitation of the FBA method is that only a limited number of environmental conditions can be considered for a specific biomass equation (or objective function) for gene essentiality.

**Table 1.** List of Features and the software packages used for feature calculation.

| Feature Types | Features name | Abbreviation of features name | # of features | Software Packages | Programming Languages |
| --- | --- | --- | --- | --- | --- |
| **Topological analysis of reactions and flux-coupled sub-networks** |  |  |  |  |  |
| Reaction Network | Degree Centrality | TF_RN_DC | 8 | The COBRA Toolbox to generate the reaction network from Genome scale metabolic network (.mat)<br>"igraph" for network analysis(42) | MATLAB, R, Perl |
|  | Eigenvector Centrality | TF_RN_EC |  |  |  |
|  | Eccentricity | TF_RN_ET |  |  |  |
|  | Hub Score | TF_RN_HS |  |  |  |
|  | Authority Score | TF_RN_AS |  |  |  |
|  | Page Rank | TF_RN_PR |  |  |  |
|  | Betweenness Centrality | TF_RN_BC |  |  |  |
|  | Number of triangle | TF_RN_NT |  |  |  |
| Flux<br>Coupled<br>Network | Degree Centrality | TF_FC_DC | 8 | F2C2 tool<br>v0.95b (Flux<br>Couple Analysis)<br>"igraph" for<br>network<br>analysis(42) | MATLAB,<br>R, Perl |
|  | Eigenvector<br>Centrality | TF_FC_EC |  |  |  |
|  | Eccentricity | TF_FC_ET |  |  |  |
|  | Hub Score | TF_FC_HS |  |  |  |
|  | Authority Score | TF_FC_AS |  |  |  |
|  | Page Rank | TF_FC_PR |  |  |  |
|  | Betweenness<br>Centrality | TF_FC_BC |  |  |  |
|  | Number of triangle | TF_FC_NT |  |  |  |
| Features derived from the coding nucleotide sequences |  |  |  |  |  |
| Derived<br>features | Nucleotide content | NS_DF_NC | 4 | In house Perl<br>script | Perl |
|  | Effective Number of<br>Codons | NS_DF_ENC | 1 | EMBOSS<br>package version<br>6.6.0-1(43) | Perl |
|  | Codon Adaptation<br>Index | NS_DF_CAI | 1 | EMBOSS<br>package version<br>6.6.0-1(43) | Perl |
| Information-theoretic<br>features | Mutual Information<br>(MI) | NS_ITF_MI | 16 | in house Perl<br>script | Perl |
|  | Conditional Mutual<br>Information (CMI) | NS_ITF_CMI | 64 | in house Perl<br>script | Perl |

| Features derived from protein sequences |  |  |  |  |  |
| --- | --- | --- | --- | --- | --- |
| Derived features | Frequencies of the twenty amino acids | PS_DF_FA | 20 | EMBOSS package version 6.6.0-1(43) | Perl |
|  | Protein length | PS_DF_PL | 1 | EMBOSS package version 6.6.0-1(43) | Perl |
|  | Paralogy based features (Paralogy score) | PS_DF_PS | 6 | BLAST [version 2.2.26] | Perl |
| Information-theoretic features | Fourier sine coefficient | PS_ITF_FSC | 70 | in house Perl script. | Perl |
|  | Fourier cosine coefficient | PS_ITF_FCC | 80 | in house Perl script. | Perl |
|  | Average Kidera Factor | PS_ITF_AKF | 10 | in house Perl script. | Perl |

Towards this, we have previously developed high performance supervised(17) and semi-supervised(18) ML-based strategies for essential gene prediction with minimal gene essentiality information which have been tested on several model organisms (Supplementary data, Table S2). The supervised ML model is built for sufficient labeled imbalanced dataset whereas the semi-supervised ML model caters to the limited labeled dataset. The pipelines consider various biological features of the genes, such as topological network features of both the genome-scale metabolic reaction network and the flux-coupled sub-networks, along with the sequence-based features that influence the gene essentiality. Consideration of these diverse features influencing gene essentiality directly provides insights into the role of a specific metabolic reaction catalyzed by a gene, estimating it to be essential. SVM classifiers performances have previously been observed to be affected by imbalanced training datasets and use of correlated or redundant features. Our supervised ML model(17) that functions in the developed web server pipeline as “Strategy 1” mitigates these shortcomings by using a SVM-based classification method which generates large number (1000 datasets) of balanced training sets ensuring that each gene is sampled at least once. In addition, the algorithm is also unique in its implementation of a Recursive Feature Elimination technique (SVM-RFE) that selects the most contributing genotype and phenotype features. As a result, our supervised ML strategy outperforms the existing supervised models for essential gene classification by predicting individual essential genes. Along with this, due to the incorporation of reaction-gene combinations, it is able to predict the associated metabolic reaction for the gene that is predicted to be essential. However, for cases of limited labeled data, ML strategy 1 performs poorly. Thus, for organisms with highly limited essentiality information, our integrative semi-supervised ML model(18) is incorporated in the web server pipeline as “Strategy 2”. The dearth of essentiality information is overcome by using a dimension reduction technique, Kamada-Kawai algorithm through LapSVM classifier that generates a distinguishing pattern between essential and non-essential genes by projection of high dimensional data onto a 2D circular layout. This results in highly accurate prediction accuracy (p <0.01) and thus significantly performs well for all organisms(18). The most distinctive feature of this semi-supervised model is that it can predict with as minimal as 1% labeled data with a statistically significant accuracy. In this strategy, an additional score SSMSS is developed for the first time, that measures the best model performance which also signifies a corresponding high auROC value. Here, we present a one-stop integrative web server platform PRESGENE at https://presgene.ncl.res.in for essential gene prediction in both prokaryotes and eukaryotes. PRESGENE is an online essential gene prediction server that hosts our previously published ML strategies(17,18) with a noteworthy capability of utilizing 289 biological features. This web server provides the user with two powerful ML-based prediction strategies that work accurately for essential gene prediction in the cases with ample as well as highly limited essential gene information.The user-friendly Graphical User Interface (GUI) of PRESGENE specially benefits biologists with limited knowledge of programming to implement ML-based prediction of essential genes in lesser studied organisms with limited experimental labeled data.

## 2 PRESGENE: Importance and Necessity

The gene essentiality information helps prioritize a set of crucial genes and their functional properties which may serve as important drug targets against various infectious diseases such as Cutaneous and Visceral Leishmaniasis, Tuberculosis, Typhoid, etc. The study of the mammalian essential genes also provides evidence for identifying important therapeutic targets and biomarkers for the treatment of cancer and other diseases. For example, in breast and ovarian cancer, homozygous BRCA 1 and BRCA 2 genes loss of function prompt the cancer cell to become dependent on poly ADP-ribose polymerase (PARP). This knowledge is exploited to treat ovarian cancer with PARP inhibitor – Olaparib(19). From the evolutionary standpoint, a distinct correlation between gene essentiality and its impact on conservation is suggested in a class or family of organisms. For instance, in *Escherichia coli*, roughly 33% of essential genes are non-essential in *Bacillus subtilis* (20). On the other hand, the study of essential genes has also been exploited in the synthetic reconstruction of the organism and in Food microbiology and industrial bioprocessing, where the essential genes and their functions in plants, animals, and microorganisms are used to produce food, biofuel, and biocatalyst at a large scale(21,22).

The essentiality of a gene varies from organism to organism, depending on the complexities of the cellular structure. To address the differences in the cellular complexities, different types of experimental protocols need to be designed(23,24). However, these techniques work well with model organisms for which a standardized protocol for gene essentiality identification is available.

Various biological features of the genes, such as topological network features of both the genome-scale metabolic reaction network and the flux-coupled sub-networks, along with the sequence-based features influence the gene essentiality. The commonly used topological network features, such as centrality measures highlight the biological significance of an enzyme in a network(25). Generally, a central and highly connected enzyme in biological networks is often essential as it represents an important hub within the network(26). If this hub node is blocked, then the whole pathway might be disrupted. Flux coupling network provides insights into the reaction subsets that are either coupled with each other via flux or represent a set of block reactions, given specific environmental exchange constraints(27,28). Consideration of these diverse features influencing gene essentiality directly provides insights into the role of a specific metabolic reaction catalyzed by a gene, deeming it to be essential.

## 3 Methods

### 3.1 Prediction Algorithms: Supervised and semi-supervised ML models

The web server provides the user with two ML strategies to choose from for their model data training and the essential gene prediction. Depending upon the availability of the labeled data for the query organism, the server is embedded with two ML algorithms:

#### 3.1.1 ML strategy 1

ML strategy 1 is developed to annotate and predict gene essentiality information for less studied organisms, where the experimentally known and labeled dataset is sufficient (≥30%) but imbalanced(17). This supervised ML strategy was trained for prokaryotes on *Escherichia coli* K12 MG1655 metabolic graph since most of the experimental data is available and the essentiality of almost all genes has been previously tested in varied environmental conditions. The training dataset for other two prokaryotes, *Brevundimonas subvibriodes* ATCC 15264 and *Helicobacter pylori* 26695 was obtained from DEG (Database of Essential Genes) v13.3(29) where experimentally labeled dataset was available. The classes considered for classification by the algorithm were ‘Essential’ with label ‘E’ for essential genes and ‘Non-essential’ with label ‘N’ for non-essential genes.

The brief steps followed in the prediction algorithm are as follows:

##### 3.1.1.1 Dataset preparation and Feature curation

The gold-standard training dataset was generated using metabolic genes from genome-scale metabolic reconstruction of model organism e.g. *Escherichia coli*. However, it is to be noted here that it is also possible to use other organisms to generate training dataset for which sufficient labeled data is available. Further, reaction-gene combinations (R_a_-G_b_) were created for network reconstruction. The total training dataset finally consisted of 4094 metabolic reaction-gene pairs. This was followed by extraction of sequence-based, gene expression-based and metabolic networks and flux-coupled network-based features that were assembled for each reaction-gene combination (Table 1). For obtaining higher classification performance, for the first time, our strategy has included network topological features from Flux Coupling Analysis(FCA)-based subnetworks that account for the inherent limitation of environmental dependence in calculation of flux distributions. FCA was performed on the iJO1366 network using F2C2 tool v0.95b(28). The training dataset was further balanced to avoid bias towards a particular class.

##### 3.1.1.2 Feature selection

SVM-RFE (Recursive Feature Elimination)(30) technique is implemented for selection of the most contributing genotype and phenotype features using WEKA version 3.8(31). Best set feature identification is performed through top ‘n’ feature combination using Sequential minimal optimization(SMO)(32) followed by 10-fold cross validation and auROC. The details of the Best Feature Combination technique (BFC) for best feature set selection can be referred to in our previous publication(17).

##### 3.1.1.3 Parameter optimization, performance metrics and model testing

Best model was identified by globally optimal hyperplane fit. A 10 fold-cross validation on 10000 datasets was performed by tuning SMO penalty parameter (*C*) and the one giving highest average auROC was selected for best feature combination. The model performance is evaluated using a weighted metric (Eq 1) with respect to the model’s classification of both class instances E and N.

Let *M* be the total set of performance metrics.

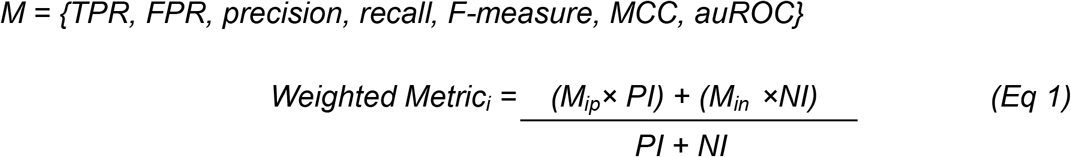

Here, *PI* is the number of positive instances, and *NI* is the number of negative instances. *M_ip_* is the performance metric for the positive class, *M_in_* is the performance metric for the negative class, where *i*∈ *M*(17).

#### 3.1.2 ML strategy 2

ML strategy 2 is developed for the prediction of gene essentiality where the experimentally known and the labeled dataset is limited (≥1%) for model training(18). A graph based semi-supervised learning method Laplacian SVM is implemented, which is based on a manifold regularization framework(33).

##### 3.1.2.1 Training dataset preparation

The two types of features as described above: topological features and sequencebased features were calculated on 12 organisms for training the semi-supervised model (Table 1). Details regarding calculation of these features can be referred to in Nandi *et al*. for details(18). In order to achieve model consistency, two types of datasets were prepared. The first type consists of 80% data points of limited labeled data for training and 20% for blind testing. The labeled data point percent is significantly varied (i% labeled from 100-i%) through randomized selection for diverse training, ensuring equal probability of Essential and Non-essential labels. In the second type, essentially for organisms where overall gene essentiality information is close to null, the whole data set (100%) is used for training purposes(18).

##### 3.1.2.2 Feature selection and dimensionality reduction

The chance of redundant features occurring is high due to the unknown contribution of the 289 features in the dataset. Thus, an unsupervised feature selection method based on the space filling concept is being applied(34). This method selects the features based on a coverage measure. This measure estimates the spatial distribution of the data points in a hypercube, thus ensuring uniform distribution of the points in a regular grid in the data space. This method does not require prior information of the output variable. Further, to reduce dimensionality, a K-Nearest Neighbour (KNN) based force-directed layout algorithm Kamada-Kawai(35) using “dimred” package in R(36). This algorithm clusters data points by minimizing the total energy. This is followed by application of the semi-supervised classifier Laplacian SVM using “RSSL” package in R(37).

##### 3.1.2.3 Performance testing and Best Model selection

It was admissible that the previously used performance metrics e.g. TPR, MCC, FPR would not be significantly applicable in a scenario of limited labeled data. Hence, a new measure called Semi-Supervised Model Selection score (SSMSS) has been proposed for the selection of the best model(18). The equation for calculation of SSMSS score is as follows:

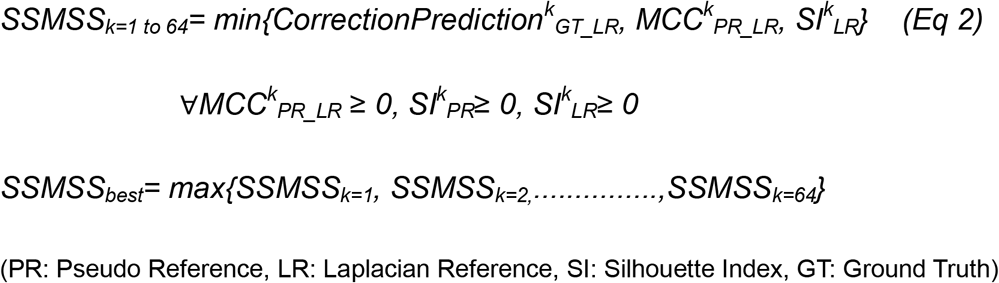

Where *k* is the *k^th^* model among 64 total models generated with a particular parametric combination and SSMSS_best_ (Eq 2) calculates the best scoring model(18). The model was further validated on the twelve organisms with well-annotated genes essentiality information that was obtained from the OGEE database(38).

#### 3.2 Features Calculation for ML Strategy 1 and ML Strategy 2

Broadly two types of features are calculated for the training and annotation of the essential genes, *viz*, the network topological features and the sequenced-based features (Supplementary data, Figure S1).The topological features of the reaction network and flux-coupled sub-network are derived from the genome-scale metabolic network of the organism. On the other hand, the sequence-based features were calculated and integrated for each reaction-gene pair based on the Gene-Protein-Reaction (GPR) rule. Integration of the diverse set of features gives insights into the specific role of the gene in the metabolic network. A total of 289 features for each reaction-gene pair can be computed to generate training and test dataset using the PRESGENE webserver. Table 1 enlists the Features and the background software packages and programming languages used for the automation of feature calculation in the PRESGENE webserver. A brief description of each of these features used for the gene essentiality prediction has been discussed in our previous work(18).

## 4 Web server architecture and Implementation

The proposed webserver has three processes, *i.e*., Training dataset Preparation, Model training, and Prediction. The workflow of the PRESGENE web server is elucidated in Figure 1A.

**Figure 1.**
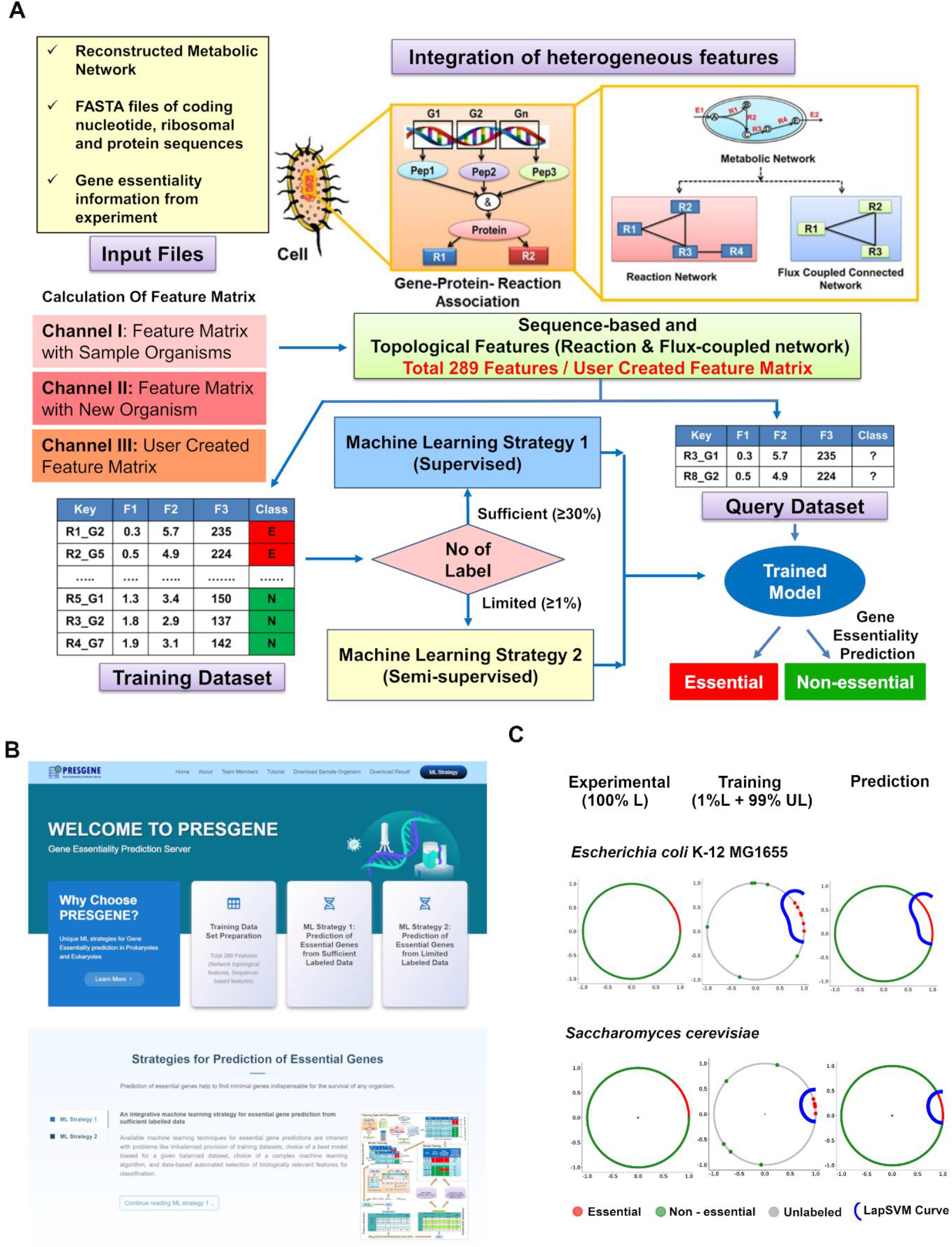
(A) Workflow for PRESGENE web server (B) Snapshot of Home page of PRESGENE web server (C) Visualization of the outcome of the Machine Learning strategy 2. Essential, non-essential, and Unlabeled reaction gene pairs are colored Red, Green, and Gray respectively. The learning curve for the best-trained model by LapSVM is colored with blue. The left circle represents the original data set with labeled data points. The middle circle shows the training data set with the learning curve, and the Right circle represents the prediction labeled with the learning curve.

### 4.1 Data input and training dataset preparation

Five input files are required for the training dataset preparation: (i) fasta file containing the coding nucleotide sequences of the genes of the organism, (ii) the ribosomal fasta file, (iii) fasta file containing the protein sequences, (iv) the genomescale reconstructed metabolic network in (*.mat) format and (v) available gene essentiality information (*i.e*., labeled data) from experiments for building the ML model. The server provides 14 sample organisms, including 9 prokaryotes and 5 Eukaryotes (**Supplementary Data, Table S2**). The fasta files of the coding nucleotide sequences, the ribosomal fasta file, and fasta files containing the protein sequences can be generally obtained from the NCBI(39) and the ENSEMBLE(40) databases. In addition, the Genome-Scale Reconstructed Metabolic Networks are available throughout the literature and the BIGG database(41). The experimental data for the gene essentiality information can be obtained from the OGEE(38), DEG(29) databases, and various experimental studies reported in the literature.

### 4.2 PRESGENE web interface and Functionality

The web interface of PRESGENE is designed in such a way that users can easily interact and navigate through the interactive web pages. The “Homepage” of the webserver contains all the necessary tabs like “About PRESGENE”, “Tutorial”, “Download Sample Organism”, “ML Strategy”, etc. The webserver homepage also provides a detailed description of the proposed machine learning strategy 1 (ML Strategy 1) and machine learning strategy 2 (ML Strategy 2) for essential gene prediction (Figure 1B). Users can perform analysis with a new dataset by providing the required input files for the calculation of the features based on their choice. Alternatively, the PRESGENE server also has a provision for the prediction of essential genes from a user-uploaded training dataset containing their own feature table. Model training can be performed using our two strategies (ML Strategy 1 and 2) depending on the availability of the labeled data.

The server provides the users with three channels or ways for predicting the essential genes via the PRESGENE server (Supplementary Data, Figure S2A). The Channel I provides the option to the user to test the pipelines on 14 sample model organisms, including both prokaryotes and eukaryotes. The user can choose to vary the percentage of labeled data to be used for the prediction of the essential genes. The results produced for these model organisms through the server can be directly incorporated by the users in their own study for prediction of drug targets or other applications. Other than these 14 sample organisms, the prediction of essential genes for a new organism using the PRESGENE server can be implemented in four simple steps. This option has been provided in the Channel II. To prepare the training dataset, the user needs to provide the name of the organism and five input files. The input files containing the GSRMN (Genome-Scale Reconstructed Metabolic Network) in (*.mat) format, fasta files of nucleotide sequence, ribosomal sequence, protein sequence, and the labeled dataset (.csv format) can be uploaded through the “Input File” navigation tab (Supplementary Data, Figure S2B). It is to be noted that all input files should maintain a uniform nomenclature for the genes. Detailed formats of these required input files have been explained in the Tutorial provided under the Tutorial tab.

Channels I and II then direct the user to the Dataset Preparation (Feature Matrix Calculation) tab to calculate and predict essential genes using the ML1 or ML2 strategies. The “Dataset Preparation (Feature Matrix Calculation)” tab allows the user to choose the set of biological features that the user wishes to consider for the gene essentiality prediction (Supplementary Data, Figure S1). However, it is recommended to consider all 289 biological features for higher accuracy and better prediction of essential genes. Through Channel III, the server additionally provides the user with an option to incorporate and test the influence of other biological features (apart from the existing 289), calculated and provided to the server in the form of a user defined Feature Matrix. This matrix forms the training dataset of the pipeline and should include the various features as columns and the reaction-gene combinations (samples) of the metabolic network as rows. The last column of the matrix should contain the gene essentiality information as E (Essential), N (Non-Essential), or UD (Undefined) as target variables. Channel III will directly take the user to the Training and Prediction tab of the ML pipeline.

### 4.3 Training and Prediction

Based on the availability of the experimentally labeled data, the user can then train the model using either ML Strategy 1 (if labeled data ≥30% of the total dataset) or ML Strategy 2 (if labeled data ≥1% of the total dataset). The performance metrics of the model are displayed on the “Training & Prediction” tab (Figure 2). In addition to the supervised performance metrics such as TPR (True Positive Rate), FPR (False Positive Rate), Precision, Recall, F-measure, the area under the receiver operating characteristic curve (auROC), accuracy, and MCC (Matthew’s correlation coefficient) in ML Strategy 1, PRESGENE offers a novel scoring technique SSMSS (Semisupervised Model Selection Score), for the section of the best model using ML Strategy 2 where the calculation of the supervised metrics is difficult. Additionally, PRESGENE allows the user to vary the feature set and recalculate the feature matrix to observe the variations in the prediction accuracy and the role of different features on gene essentiality prediction.

**Figure 2.**
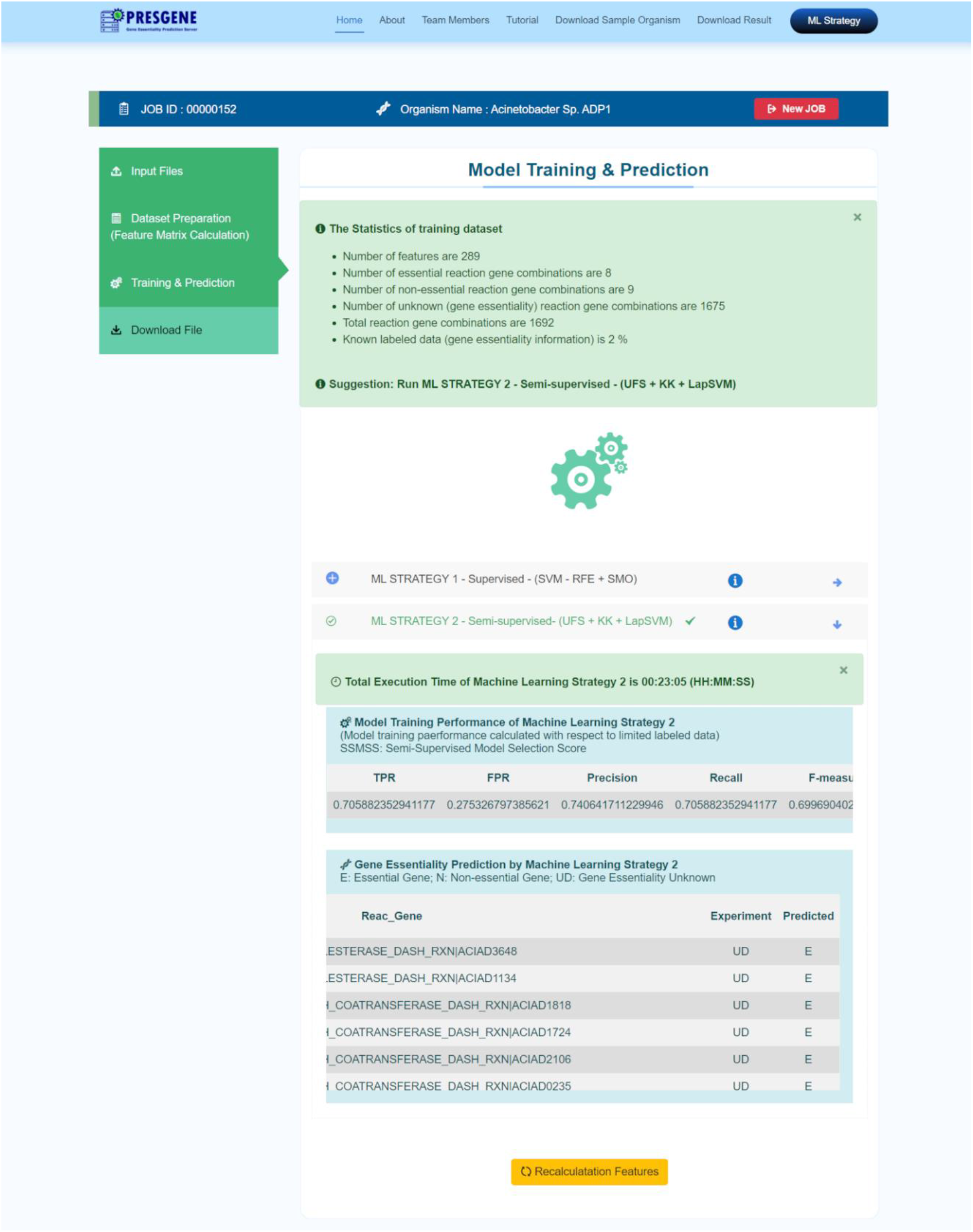
Training and Prediction tab for visualizing the prediction outcome. The ML Strategy pipeline Result Tab “Model Training and Prediction” displaying model Training performance Metrics such as TPR, FPR, Precision and the predicted essential genes list as the final result output.

The results along with the calculated feature table generated for the prediction of the essential genes, using the PRESGENE server, can be downloaded in.csv format by the user from the “Download File” tab. The results will be available for 15 days in the server and during this period it can be retrieved anytime from the Download Result tab using the JOB ID. A detailed Tutorial has been provided for the benefit of the users.

### 4.4 Prediction efficacy and performance

The performance of PRESGENE was assessed based on the training and prediction accuracy as well as the universality of the proposed supervised model strategy, ML Strategy 1. A comparative performance testing of ML Strategy 1 with a previously established model proposed by Hwang *et al.(8*) was carried out. The Hwang *et al*. strategy uses sequential minimal optimization (SMO)(32) algorithm and a linear kernel-based SVM, whereas our model implements SVM-RFE (Recursive Feature Elimination) technique. The curated dataset of model organisms from our study as well as the dataset used in Hwang *et al*. study was used for the performance testing. The comparison was quantified by performance measure metrics (*i.e*. Precision, Recall, F-measure and MCC) (Table 2). In terms of the training accuracy, our model shows significantly improved classification performance as can be observed from Table 1 with an improved MCC and F-measure values for both Hwang *et al*. dataset as well as our curated dataset. For example, our strategy produced a F-measure of 0.826, a significant increase from the F-measure of 0.784 by Hwang’s strategy. In the case of Strategy 2, the improved semi-supervised ML-based algorithm, 1% labeled data of the twelve organisms was used. To compare the performance of our classifier with existing classifiers, different supervised classifiers like Random forest (RF), Naive Bayes (NB), Logistic regression (LR) and decision tree (DT) were used for testing (Figure 3). Our Laplacian SVM based classifier was found to outperform all other methods significantly. The semi-supervised strategy in the server has performed with equal accuracy in the case organisms *Leishmania donovani* and *Leishmania major, which* has been demonstrated previously in details as a case study(18).

**Figure 3.**
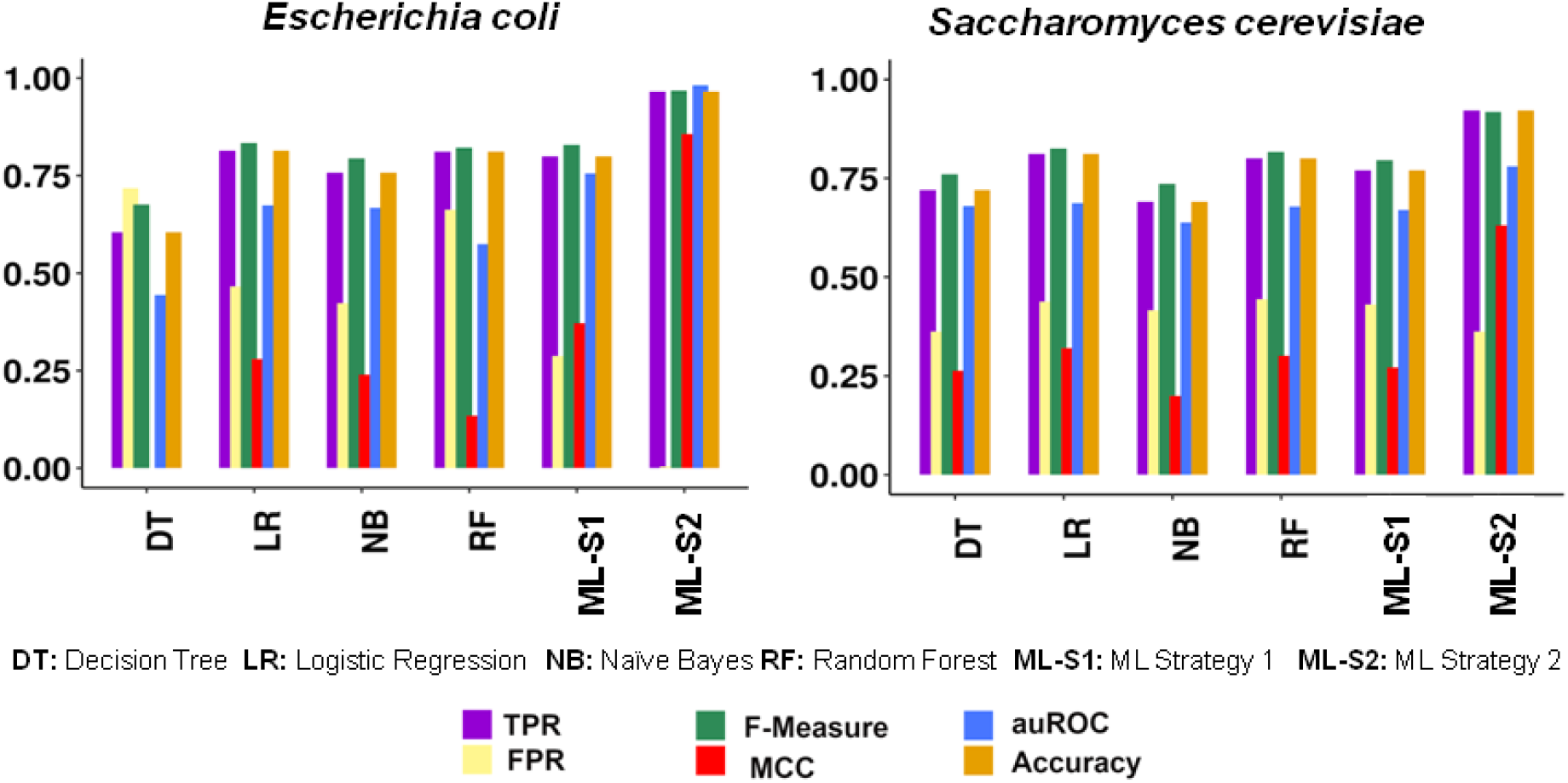
Comparison of the predictive performance of the ML strategies of PRESGENE with other supervised methods. Comparison of the performance of ML Strategy 1 and ML Strategy 2 used by PRESGENE with other available supervised classifiers i.e., Decision Tree (DT), Logistic regression (LR), Naive Bayes (NB), Random Forest (RF) based on 1% labeled data for the two case study organisms, prokaryote: Escherichia coli and eukaryote: Saccharomyces cerevisiae. The X-axis represents the different types of performance metrics for machine learning strategies; Y-axis represents the value of performance metrics.

**Table 2.**
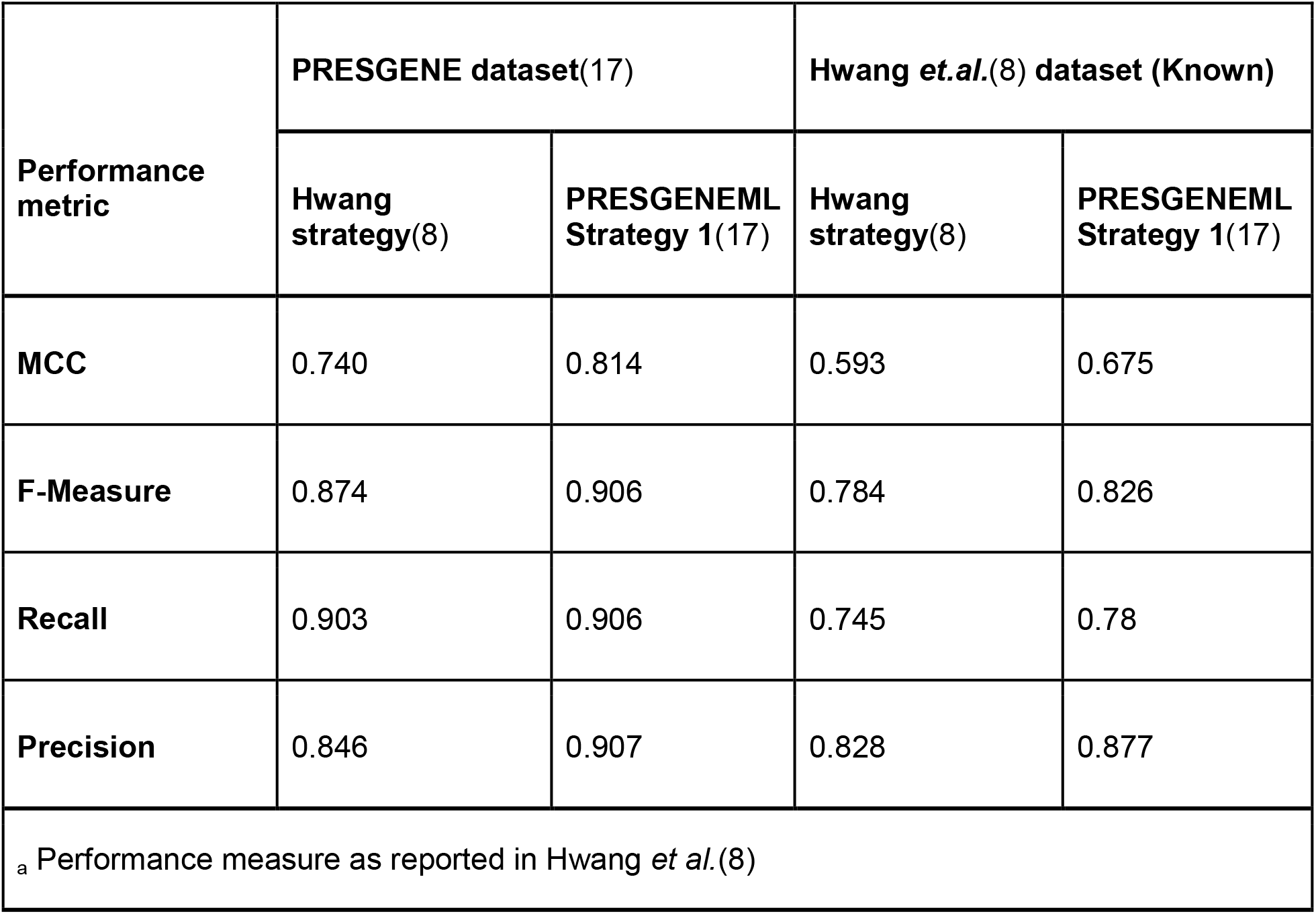
Performance testing of ML Strategy 1(17) i.e., supervised strategy by comparison with Hwang *et al*. study(8).

### 4.5 Case Study to assess Performance efficacy

The server hosts a total of 14 model organisms for which the ML models are trained with 1% labeled data. The prediction performance is assessed by SSMSS score as well as the semi-supervised metrics (i.e., TPR, FPR, F-measure, MCC, auROC, and accuracy)(18). For example, the outcome of the ML strategy 2 for prokaryotic model organism *Escherichia coli* and eukaryotic model organism *Saccharomyces cerevisiae* can be visualized as three circles (Figure 1C). The first circle represents the circular projection of the whole data set in 2-D with gene essentiality information from the experiment. The second circle shows the training data set with 1% labelled & 99% unlabeled data. The third circle shows the predicted gene essentiality label from the best-trained model and the LapSVM curve based on SSMSS scoring for the best trained model. It is observed that the ML strategy 2 model performed well for both organisms (similar circular patterns from experiment and prediction). Similarly, for the same case study organisms *Escherichia coli* and *Saccharomyces cerevisiae*, the comparison of performances of our PRESGENE ML Strategy 1 and ML Strategy 2 with other supervised methods (as mentioned in section 4.4) are observed to be significantly higher, with ML Strategy 2 achieving an Accuracy value of 0.899 for *Escherichia coli* and 0.921 for *Saccharomyces cerevisiae* (Figure 3).

### 4.6 User Interface Design

Bootstrap 4 framework has been used for designing the front end of the server. The programming languages MATLAB, Perl, R, and PHP have been used to write code for the automation of feature calculation and deployment of the machine learning pipelines (ML Strategy 1 and ML Strategy 2) for the essential gene prediction. The present configuration of the PRESGENE server is Intel(R) Xeon(R) CPU E5-2680 @ 2.70GHz with 32 CPUs and 128 GB RAM. This allows maximum four to five users to use the PRESGENE web service simultaneously. In future, the hardware configuration of the server can be upgraded to accommodate a greater number of users simultaneously.

A key limitation of the server lies in the fact that both the ML strategies fail to execute if the genome-scale reconstructed metabolic network of the organism and a minimum of 1% labeled dataset are not available. For time being, one can give the reconstructed metabolic network in MAT-file (.mat) format to make it comprehensive for the web-server to process it further. Further, we are working to incorporate additional data formats of genome-scale models so that one will be able to use automated GSMs from a different source in the near future.

## 5 Conclusion and Impact of PRESGENE

In this paper we explained in detail the architecture of PRESGENE web server that implements our previously introduced ML strategies (17, 18) for essential gene prediction. This web server is intended to be used by biologists for prediction of essential genes in novel prokaryotic and eukaryotic organisms which can influence better characterization of novel organisms.

The main impact of our web server lies in its ability to seamlessly classify essential and non-essential genes implementing our supervised and semi supervised ML algorithms for organisms with extremely limited essential gene information, such as in cases of up to only 1% labeled data from the organism’s genome. Further, the algorithm uses a vastly diverse set of features (stems from FBA sub network, metabolic gene-reaction pair), which has previously not been implemented that improves the prediction accuracy manifold in organisms with least known essentiality data. The supervised ML strategy mitigates the inherent problems with unbalanced training datasets, feature bias with its unique implementation of SVM-RFE technique with higher classification performance and has the ability to capture a minimal set of essential genes that contribute to essentiality. On the other hand, the semisupervised ML Strategy excels in its performance for prediction on highly limited essentiality information for unknown organisms by combining LapSVM classifier for training along with Kamada-Kawai dimension reduction technique and also presents a prediction accuracy monitoring score SSMSS for the proposed technique. These high performance prediction algorithms benefit a wide variety of users. Additional advantages of using our web server include: 1) One can implement our ML strategies on the 14 model organisms for which the entire required data is provided within the server; 2) One can use the server for their organism of interest with option to choose either of the ML strategy based on the availability of labeled data; 3) One can easily explore from a plethora of features (currently available, 289) for training set preparation and can also customize the feature matrix; 4) A detailed tutorial guides the user step-by-step process for a seamless use of web server with just click-based operation and thus can be used by any biologists with limited or no knowledge of computational methods.

Hence, PRESGENE will be invaluable to experimental and computational biologists by providing a well-validated and standardized platform to annotate gene essentiality of less-explored organisms with minimal information on labeled data. The essential genes predicted using the platform have broad applicability and will help identify novel therapeutic targets against disease-causing organisms for antibiotic and vaccine development.

## Abbreviations

ML: Machine Learning
FBA: Flux Balance Analysis
KNN: K-Nearest Neighbour
SVM: Support Vector Machine
FCA: Flux Coupling Analysis
RFE: Recursive Feature Elimination
BRCA: Breast Cancer
SMO: Sequential minimal optimization
TPR: True Positive Rate
FPR: False Positive Rate
ROC: Receiver Operating Curve
MCC: Matthews Correlation Coefficient
NCBI: National Centre for Biotechnology Information

## 6 Acknowledgment

The authors acknowledge Mr. Arun Kumar Kundu from St. Xavier’s College, Kolkata for helping in front end design of the webserver and Mr. AnirudhMurali for maintenance of the web server.

## 7 Author contribution

**Sutanu Nandi:** Data curation, Methodology, Web server Development, Writing - Original Draft **Gauri Panditrao:** Data Analysis, Validation, Web server Implementation and Visualization, Writing -Original Draft **Piyali Ganguli**: Implementation and Visualization, Writing-Original draft, Review and Editing **Ram Rup Sarkar**: Conceptualization, Investigation, Supervision, Writing-Review and Editing.

