## Supplementary data for "PRESGENE: A web server for PRediction of ESsential GENE using integrative machine learning strategies"

**Table S1. Comparison of different machine learning methods for predicting essential genes.**

| **Organisms** | **Biological features** | **Feature Selection** | **Dimension Reduction** | **ML Classifier** | **Availability of Source Code /Webserver** | **Availability of Dataset** | **References** |
| --- | --- | --- | --- | --- | --- | --- | --- |
| *S. cerevisiae* | PPI, Sequence | No | No | Neural network and SVM | No | No | (1) |
| *E. coli and*  *S. cerevisiae* | PPI, Sequence | Yes | No | Bayesian algorithm | No | Yes | (2) |
| *S. cerevisiae* | Sequence | No | No | Bayesian algorithm | No | No | (3) |
| *S. cerevisiae* | PPI, Sequence | Yes | No | *k*-Nearest neighbor, SVM | No | No | (4) |
| *E. coli* | PPI | No | No | Decision tree | No | No | (5) |
| *E. coli* | MN, Sequence | Yes | No | SVM | No | No | (6) |
| *S. cerevisiae* | PPI and others | No | No | Decision tree | No | No | (7) |
| Multiple organisms | MN, Sequence | No | No | Decision trees and SVM | No | No | (8) |
| Multiple organisms | PPI, Sequence | No | No | Bayesian, logistical regression, decision tree and CN2 rule | No | No | (9) |
| *Mus musculus* | PPI, Sequence | No | No | Logistic regression, random forest and SVM | No | No | (10) |
| Multiple organisms | PPI, Sequence | No | No | Bayesian algorithm | yes | No | (11) |
| *S. cerevisiae* | PPI | No | NO | Logistic regression | No | No | (12) |
| Multiple organisms | PPI, Sequence | No | No | Bayesian algorithm | No | No | (13) |
| *Aspergillus fumigatus* and yeast | Gene Co expression Network, Sequence | No | No | Bayesian algorithm, logistical regression, decision tree and CN2 rule | No | No | (14) |
| Multiple organisms | Sequence | No | No | SVM | No | No | (15) |
| *Homo sapiens* | PPI, Sequence | No | No | SVM, logistic regression and decision tree | No | No | (16) |
| *Arabidopsis thaliana*, *Oryza sativa* and  *S. cerevisiae* | PPI, Sequence | No | No | Random forest | No | No | (17) |
| Multiple organisms | Sequence | No | No | SVM | No | No | (18) |
| *E. coli*, *Streptococcus pneumoniae* TIGR4 | PPI, Sequence | No | No | Principal component regression | No | No | (19) |
| *Homo sapiens* | Sequence | Yes (SVM-RFE) | No | SVM | No | No | (20) |
| Multiple organisms | Sequence | No | No | SVM | No | No | (21) |
| Multiple organisms | Sequence | Yes (LASSO) | No | SVM | No | No | (22) |
| *Drosophila melanogaster* | Sequence, PPI Network | Yes (LASSO) | No | GLM, SVM, RF, ANN | No | No | (23) |
| Multiple organisms (microbes) | Sequence | No | No | deep neural network (DNN) | Yes | Yes | (24) |

**Table S2. Organisms considered as sample model organisms to test PRESGENE server.**

| **Organism Name** | **Input files** | | |
| --- | --- | --- | --- |
|  | **FASTA files of coding nucleotide ribosomal and protein sequence**  **(RefSeq assembly accession)** | **Genome-Scale Reconstructed**  **Metabolic Network** | **Gene Essentiality Information** |
| **Prokaryotes** | | | |
| *Acinetobacter sp*. ADP1 | GCF_000046845.1_ASM4684v1 | iAbaylyiv4 (25) | OGEE (26) |
| *Bacillus subtilis subsp. subtilis str.* 168 | GCF_000009045.1_ASM904v1 | iYO844 (27) |  |
| *Escherichia coli* K-12 MG1655 | GCF_000005845.2_ASM584v2 | iJO1366 (28) |  |
| *Helicobacter pylori* | GCF_000008525.1_ASM852v1 | iIT341 (29) |  |
| *Mycobacterium tuberculosis* H37Rv | GCF_000195955.2_ASM19595v2 | iNJ661 (30) |  |
| *Pseudomonas aeruginosa* PAO1 | GCF_000006765.1_ASM676v1 | iPae1146 (31) |  |
| *Pseudomonas aeruginosa* UCBPP-PA14 | GCF_000014625.1_ASM1462v1 | iPau1129 (31) |  |
| *Salmonella enterica subsp. enterica serovar Typhimurium* LT2 | GCF_000006945.2_ASM694v2 | STM_v1_0 (32) |  |
| *Staphylococcus aureus subsp. aureus* NCTC 8325 | GCF_000013425.1_ASM1342v1 | BMID000000141098 (33) |  |
| **Eukaryotes** | | | |
| *Saccharomyces cerevisiae* | GCF_000146045.2_R64 | iMM904 (34) | OGEE (26) |
| *Caenorhabditis elegans* | GCF_000002985.6_WBcel235 | iCEL1273 (35) |  |
| *Mus musculus* | GCF_000001635.26_GRCm38.p6 | iMM1415 (36) |  |
| *Leishmania donovani* | TriTrypDB-36 | iMS604 (37) | (38) |
| *Leishmania major* | TriTrypDB-36 | iAC560 (39) |  |

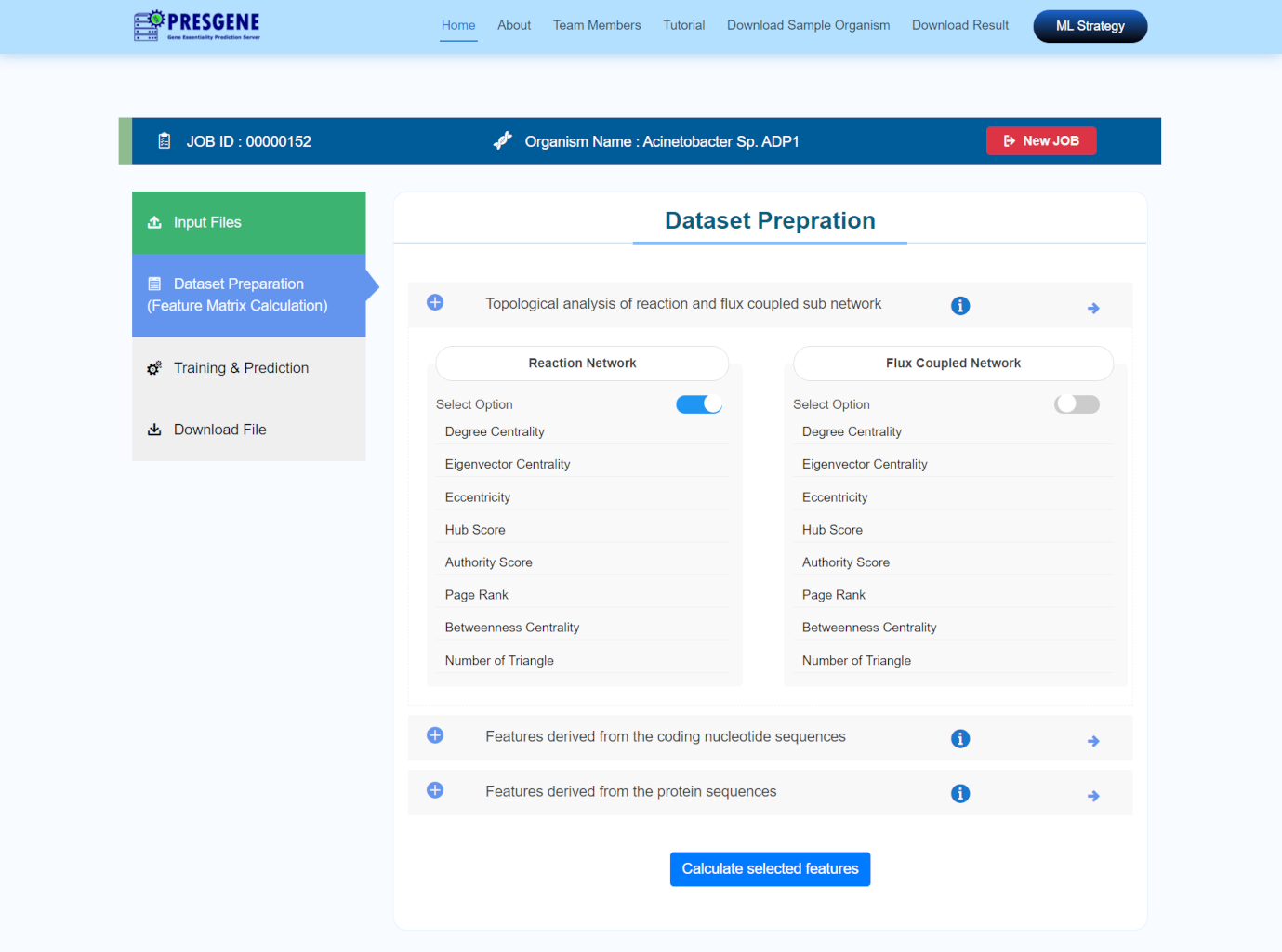

**Figure S1 Dataset Preparation (Feature Matrix Calculation) tab for feature matrix calculation.** This tab provides the user option to select the desired features for training dataset preparation

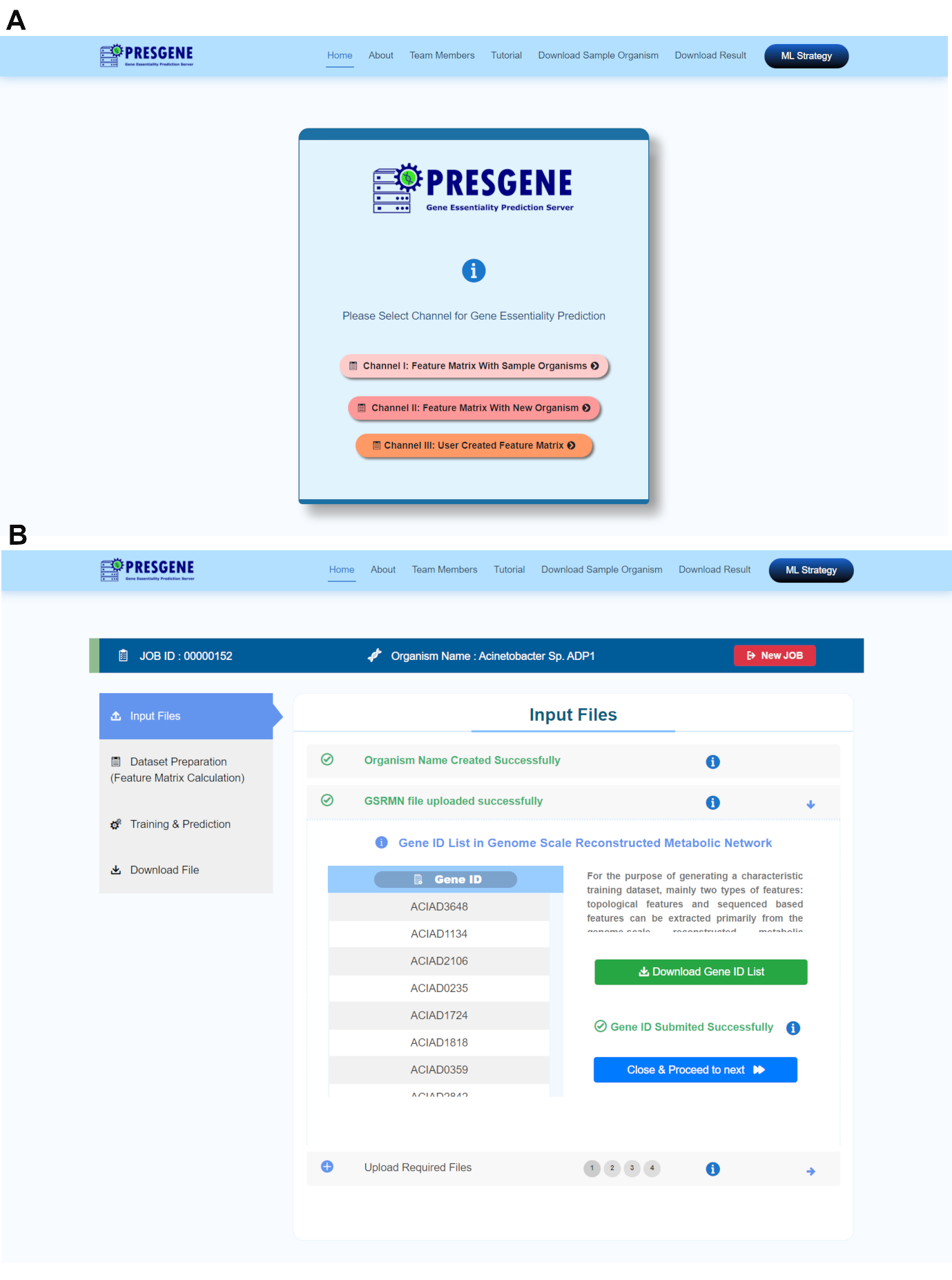

**Figure S2 ML strategies for genes essentiality prediction. (A)** The ML strategy tab allows the user to select from three channels for calculating the feature matrix **(B)** Input Files navigation tab for new organisms.
